# Jagged1 overexpression on T cells induces thymic regulatory T cells leading to thymic involution

**DOI:** 10.1101/2022.08.25.504005

**Authors:** Joanna S. Kritikou, Irene Sánchez-Pascual, Juan Pedro Muñoz-Miranda, Neha Vashist, Arnika K. Wagner, Xing-Mei Zhang, Erika Assarsson, Margarita Dominguez-Villar, Hideo Yagita, Francisco Garcia-Cozar, Peggy Riese, Robert A. Harris, Hans-Gustaf Ljunggren, Benedict J. Chambers

## Abstract

We previously described a mouse model, tg66, with a severe defect resulting in diminished thymic size and complete involution by early adulthood. In the current study, we identified overexpression of Jagged1 as a mechanism for the alterations in thymic development in tg66 mice. T cells in the tg66 thymus were skewed towards CD8^+^ T cells, and within the CD4^+^ T cell compartment there was an over-representation of Foxp3^+^ cells. Regulatory Foxp3^+^ T cells (Tregs) isolated from tg66 mice had increased ST2 and CD103 expression. These Tregs could suppress proliferation to the same extent as conventional Tregs. Corroborating these results, tg66 mice were resistant to experimental induction of neuroinflammation in a common animal model for multiple sclerosis (EAE). Using bone marrow chimeras, we recorded a stark reduction in the number of thymocytes and a corresponding increase in Tregs in the thymus of mice receiving tg66 bone marrow. Conversely, through blocking Jagged1, the number of thymocytes was significantly increased, being concomitantly associated with a drop in the frequency of Tregs. We conclude that Tregs may play a role in thymic involution and could explain early thymic involution or loss as it is observed in diseases of thymic atrophy such as Down syndrome.

## Introduction

T cell selection is conducted in the thymus via epithelial cells that present self-antigens to developing T cells (1). These thymic epithelial cells (TEC) can be divided into two groups, cortical (cTEC) and medullary (mTEC) cells. cTECs impose positive selection while, with the help of accessory cells such as dendritic cells, mTECs are involved in negative selection (1). Incoming T cells from the bone marrow are first selected on cTECs, evolving from the DN1 (CD4^−^CD8^−^CD44^+^CD25^−^) stage to DN2 (CD44^+^CD25^+^), DN3 (CD44^−^CD25^+^), and DN4 (CD44^−^CD25^−^) stages. During stages DN2 and DN3, VDJ rearrangement of the T cell receptor (TCR) begins and by the end of DN4 the cells express a TCR β chain and both CD4 and CD8. These so-called double positive (DP) T cells then rearrange their TCR α chain and with their fully formed αβTCR can bind to MHC molecules. T cells with low avidity for self-peptides will survive, proliferate, and retain either CD4 or CD8, depending on their ability to bind to MHC class II or class I, respectively. These single positive (SP) cells can then exit the thymus as immature, naïve T cells. CD4^+^Foxp3^+^ T regulatory cells (Tregs) can develop both in the thymus and the periphery, but the majority arises in the thymus as a mature, suppressive T cell subpopulation. Like effector T cells, Tregs also undergo selection in the thymus. Co-stimulatory molecules such as CD28, CD80/86 (B7), CD40 and IL-2Rβ appear to be critical for Treg development (2). Additionally, TGFβ is required to maintain Foxp3 expression and the suppressive function of peripheral Tregs (3).

Signaling through Notch receptors is evolutionarily conserved and regulates cell fate decisions during development. Four Notch genes (*Notch1-4*) and two ligand groups exist in mammals. The ligands are Jagged (Jagged1 and Jagged2) and Delta-like (DLL1, DLL3, and DLL4). Notch signaling has also been implicated in T cell development in the thymus, with the pathway enabling terminal differentiation of both CD4^+^ and CD8^+^ T cells by activating relevant transcription factors for these subtypes (4). Conversely, Notch signaling appears to be inhibitory in Tregs, since knocking out *Notch1* in T cells enhances their suppressive capacity (5). Jagged1 over-expression has the opposite effect, promoting the regulatory capacity of T cells and maintaining Tregs *in vivo* (6-8). In contrast, blocking Jagged1 exacerbates experimental autoimmune encephalomyelitis (EAE), a mouse model for multiple sclerosis (9).

While thymic involution in humans normally occurs after puberty, early involution is observed in pathological cases induced by defective genes or environmental factors (10, 11). Premature thymic involution is observed in Down syndrome (12) as well as in mouse models of the disease (13). In several other diseases such as DiGeorge’s syndrome, severe combined immunodeficiency, and Wiskott-Aldrich syndrome, thymic involution is thought to contribute to the associated decreased T cell output and symptoms (14). While it is currently not understood why the thymus undergoes shrinkage in non-pathological conditions (11), energy conservation and prevention of both lymphoid leukemia and autoimmune diseases have been proposed, although the exact mechanisms remain obscure. One factor known to be involved in thymic involution is TGFβ signaling (15, 16), which is active in TECs that support thymopoiesis. Eosinophilia has also been associated with thymic involution (17), with eosinophils expressing TGFβ and their infiltration to the thymus coinciding with the commitment to involution.

We have previously described a mouse model, tg66, which exhibits normal thymic size at birth but which rapidly shrinks with adulthood (18). In homozygous tg66 mice, no detectable thymus is evident in the mice after 28 days. Herein, we report that thymic involution in tg66 mice is due to an increased apoptotic response of their TECs. In parallel, we observed significant expansion of the Treg compartment likely resulting from Jagged1 overexpression in these mice. This increase in Treg numbers was also associated with the tg66 mice not developing symptoms in an EAE mouse model.

## Materials and Methods

### Animals

C57Bl/6 (B6), tg66, tg71 (18) and C57Bl/6 perforin (B6.*pfp*)^−/−^ (19) mice were bred and housed under standard conditions at the Department Microbiology and Tumor and Cell Biology, Karolinska Institutet, Stockholm. Tg71 are used as controls to tg66 mice, as these have a successful insertion of the transgene NK1.1, without disrupting control of JAG1 expression and with no signs of thymic disfunction. RAG2^−/−^ x common-γ chain^−/−^ (RAG2γc^−/−^) mice were maintained in IVC cages. All procedures were performed under both institutional and national guidelines (Ethical numbers from Stockholm County Council N147/15).

### Flow cytometry

All antibodies were purchased from Becton Dickenson (San Diego, CA), Bioscience (San Diego CA) or Biolegend (San Diego CA); anti-CD3 (clone 145.2C11, BD Biosciences), -Jagged1 (HMJ1-29), -NKp46 (29A1.4), -NK1.1 (PK136), -CD103 (2E7), FoxP3 (MF14), -ST2 (DIH4), -KLRG1 (2F1/KLRG1), GITR (DTA-1) -ICOS (7E.17G9) and -Siglec F (1RNM44N). Anti-granzyme C antibody was a kind gift from Timothy Ley University of Washington St Louis (20). For intracellular staining, cells were fixed in Foxp3 fixation and permeabilization buffer (eBioscience). Flow cytometry was performed using a FACSort or FACScalibur flow cytometer (Becton Dickinson, Mountain View, CA) and a CyAN ADP LX 9-colour flow cytometer (Beckman Coulter, Pasadena, CA). Data were analyzed using FlowJo software (Tree Star Inc, OR). Gating strategy for thymocytes is shown in Supplemental Fig. 1.

### RNA preparation and RT-PCR

Single cell suspensions were made of spleens from tg71 (control) and tg66 mice. Red blood cells were lysed and pellets were resuspended in phosphate-buffered saline (PBS). Live cells were counted, washed twice in PBS and kept at −20°C in RLT buffer containing 1:100 v/v β-mercaptoethanol. The rest of the extraction was performed using the RNeasy Minikit (Qiagen, Hilden, Germany) according to the manufacturer’s recommendations. Reverse transcription was performed using 1 mM oligodT and Superscript^TM^ HH RNase H-reverse transcriptase (Invitrogen Life Technologies) according to the manufacturer’s instructions. Finally, quantitative PCR was performed using a LightCycler System (Roche diagnostics) and SYBR Green PCR kit from Roche Diagnostics. The cDNA input for each population was normalized to obtain equivalent signals with Splicing. HPRT was used as housekeeping gene. The *Jag1* forward and reverse primers for the qPCR were as follows:

Forward: 5’ GCAAGACTTGTCAGTTAGATG 3’

Reverse: 5’ CTGGCAATCAGATTCTTACAG 3’

### Generation of bone marrow chimeras

Six- to eight-week old RAG2γc^−/−^ mice were sub-lethally irradiated with 500 rad followed by an additional 600 rad after 8h and were then injected with 10^6^ bone marrow cells (from tg66 or tg71 mice) in the tail vein. Donor mice were depleted of Tregs 24h before injection using the anti-CD25 antibody PC61 (BioXCell, Lebanon NH). For continued depletion of Tregs, mice were injected once a week for the duration of the study. For blockade of Jagged1, anti-Jagged1 (clone HMJI-29) antibody (9) was injected intraperitoneally (i.p.) every 3 days for the course of the study at a dose of 500μg/mouse.

### In vitro T cell suppression assay

CD8^+^ T cells were positively sorted with beads from the spleens of C57BL/6 mice (Miltenyi Biotec). The T cells were labelled with CFSE (Invitrogen Molecular Probes), washed and added to U-bottom plate at a concentration of 10^5^ cells/well in RPMI media containing 10% FBS. with anti-CD3/CD28 beads and different ratios of Tregs from tg66 or tg71 mice were added into the cultures. After 72 hours, proliferation was examined by flow cytometry.

### Induction of EAE

Recombinant protein corresponding to the N-terminal sequence of mouse myelin oligodendrocyte glycoprotein (MOG) (amino acids 1–125) was expressed in *Escherichia coli* and purified to homogeneity by chelate chromatography as previously described (21). Purified MOG dissolved in 6 M urea was dialyzed against sodium acetate buffer (10 mM, pH 3.0) to obtain a soluble preparation that was stored frozen at −20°C. Mice were anesthetized with isoflurane (Forane; Abbott Laboratories, Abbot Park, IL) and injected subcutaneously at the dorsal tail base with 100μL inoculum containing 50 μg of MOG in PBS emulsified in Complete Freund’s Adjuvant (CFA) containing 100 μg heat-killed *Mycobacterium* tuberculosis H37Ra (Difco Laboratory, Detroit, MI). Mice were weighed and scored daily for clinical signs of EAE using a 6-point scale as follows: 0, no clinical signs of EAE; 1, tail weakness or tail paralysis; 2, hindlimb paraparesis or hemiparesis; 3, hindlimb paralysis or hemiparalysis; 4, tetraplegia or moribund; and 5, death.

### Statistical Analysis

All statistical analysis was performed using GraphPad Prism software (La Jolla, CA).

## Results

### Jagged1 overexpression is associated with thymic loss

Heterozygous tg66 transgenic mice have an insertion in the NK1.1 gene under the control of the CD2 promoter that leads to loss of their thymus during adult development (18). Homozygous tg66 mice lose their thymus more rapidly than control wt or tg71 mice and the thymus is not detected after 28 days (Fig. 1A). Not surprisingly, loss of the thymus is related to a reduced number of T cells in the periphery (Fig. 1B, (18)). We found that the defect was associated with Jagged1 expression on T cells since we could find increased levels of *Jag1* at the RNA level and significant increase in the surface expression of Jagged1 on T cells in the periphery and in the thymus (Fig. 1C-F). Since CD2 is found on other immune cells, we evaluated the level of surface expression of Jagged1 also on NK cells, B cells and dendritic cells which can express Jagged1 under normal physiological conditions (Fig. 1G). While we consistently saw increased expression of Jagged1on all these cell types from the spleens of tg66 mice compared to tg71, these increases were not significantly different.

**Figure 1.**
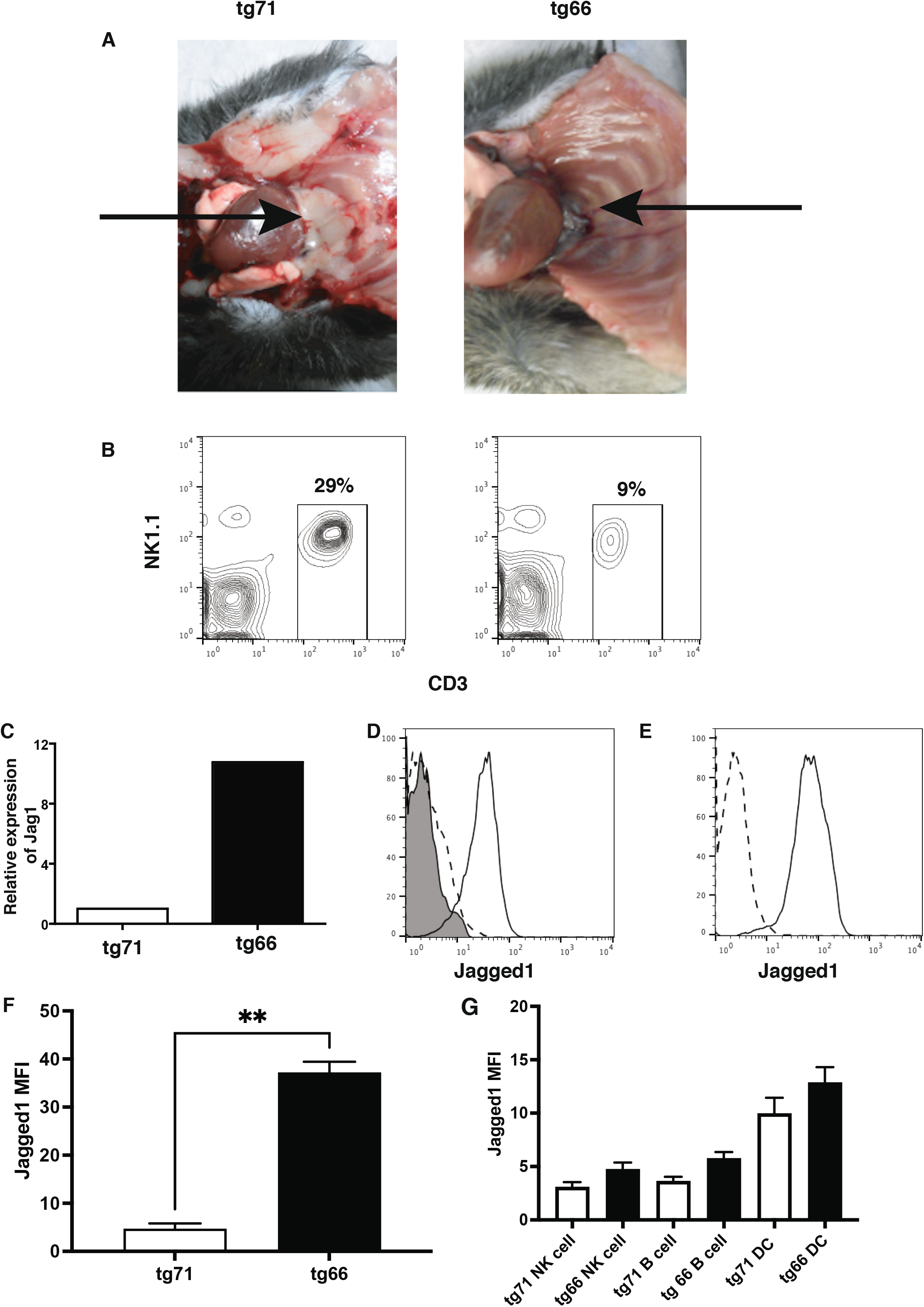
Expression of Jagged1 in tg66 mice. (A) Loss of thymus in tg66 mice. Arrows indicate position of the thymus. (B) The reduced CD3^+^ T cell population in the spleen of tg66 mice. (C) Expression of *Jag1* mRNA in T cells from tg71 and tg66 mice. (D) Surface Jagged1 expression on splenic T cells from tg66 (solid line) and tg71 mice (dashed lines). (E) Surface Jagged1 expression on thymic T cells from tg66 (solid line) and tg71 mice (dashed lines) and (F) bar graph of the expression levels comparing tg66 and tg71 (**p<0.01 Mann Whitney test n=6 mice) (G) Expression of Jagged1 on NK cells, B cells and dendritic cells from the spleen.

### Increased apoptosis of thymic epithelium from tg66 mice

We previously reported that the thymic cytoarchitecture of tg66 mice was extremely disturbed, with poor differentiation of cortical and medullary areas (18). This was reminiscent of the observation in the thymi of mice expressing Jagged1 under the *lck* promoter (22). In the latter study, a high degree of apoptosis amongst the TECs was observed, suggesting that altered T cell regulation might drive TEC cell death. We therefore examined whether this was similar in the tg66 mice. For these studies, we took day14 old mice since the thymus size was equivalent to tg71 mice. We found that there was increased frequency of apoptosis in the epithelial cells of thymi from tg66 mice (Fig. 2A-D). These data and the previously published data together suggested that Jagged1 over-expression on thymic T cells could cause alterations in these cells that led to the apoptosis and/or necrosis of epithelial cells, potentially through interacting with Notch receptors on the stroma/epithelium with Notch ligand Jagged1 on T cells.

**Figure 2.**
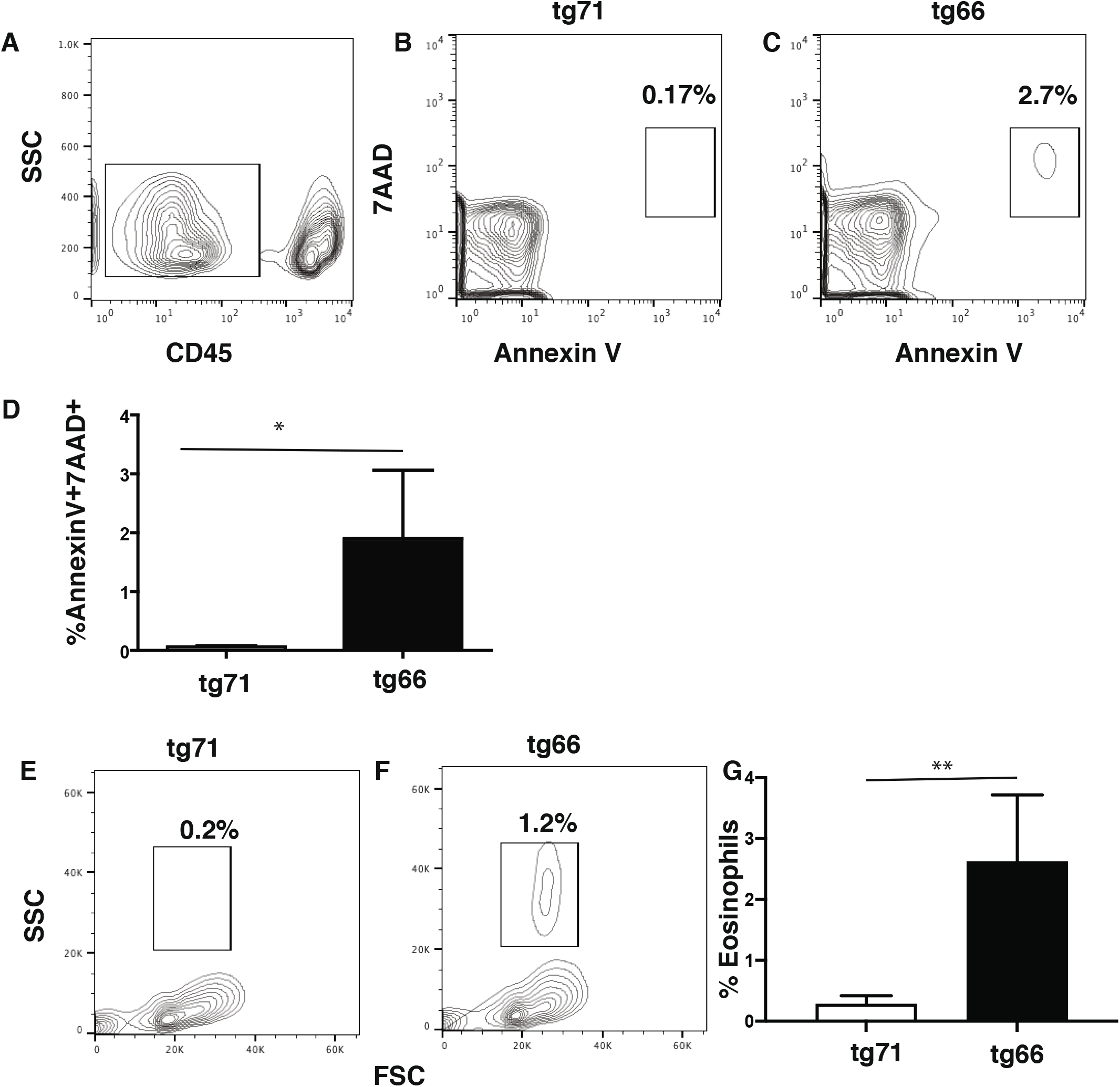
Increased apoptosis of thymic epithelial cells and eosinophil infiltration in the thymus of tg66. (A) Gating of CD45^−^ cells in the thymus of 14 day old mice (B) Apoptosis of CD45^−^ cells in the thymus of 14 days old tg71 mice. (C) Apoptosis of CD45^−^ cells in the thymus of 14 day old tg66 mice. (D) Bar graph representing the percentage of Annexin V^+^/7AAD^+^ in the thymi of 14 day old tg71 and tg66 mice. (*p<0.05 Mann Whitney test, n=4-5 mice). (E and F) Forward and side scatter plots of the thymocytes from (E) 14 day old tg71 and (F) tg66 mice after gating away dead cells. (G) Bar graphs represent the percentage of eosinophils in the thymi of tg71 and tg66 mice. (**p<0.01 Mann-Whitney test, n=5 mice)

Eosinophilic infiltration of the thymus has also been associated with thymic involution (17). When thymi from 14 day-old tg66 and tg71 mice were compared, there was a distinct increase in granulocytes observed in the thymus of tg66 mice (Fig. 2E-G). These granulocytes were primarily SiglecF^+^, indicating they were eosinophils (Supplemental Figure 1). Infiltration of eosinophils thus appears to be associated with thymic involution in tg66.

### Jagged1 over-expression increases frequency of Tregs in tg66 mice

We had previously published that T cell development in the tg66 mice was skewed towards CD8^+^ T cells with a marked reduction in CD4 T cells (Supplemental Figure 2)(18). However, since Jagged1 also plays a role in CD4^+^ Treg cell development (8, 23-26), we examined the expression of Foxp3 in the remaining CD4^+^ T cell population from the spleens of adult mice 7-8 weeks old. In control tg71 mice, the frequency of Foxp3^+^ T cells was approximately 15% of the CD4 T cells, while in the tg66 mice the frequency was significantly higher at around 30% (Fig. 3A). Upon examining the thymus of 14 day-old tg66 mice, approximately 20% of the CD3^+^CD4^+^ SP T cells were Foxp3^+^, compared to approximately only 5% in tg71 mice (Fig. 3B) suggesting a developmental skewing of CD4^+^ T cells to Tregs in the tg66 mice. To test if the Tregs generated in tg66 mice were functional, we performed proliferation suppression assays in which CD8^+^ T cells were stimulated with anti-CD3/CD28 beads and different ratios of splenic Tregs from tg66 or tg71 mice were added into the cultures. The proliferation rates of CD8^+^ T cells were almost identical, suggesting that the Tregs from the tg66 mice were functional (Fig. 3C).

**Figure 3.**
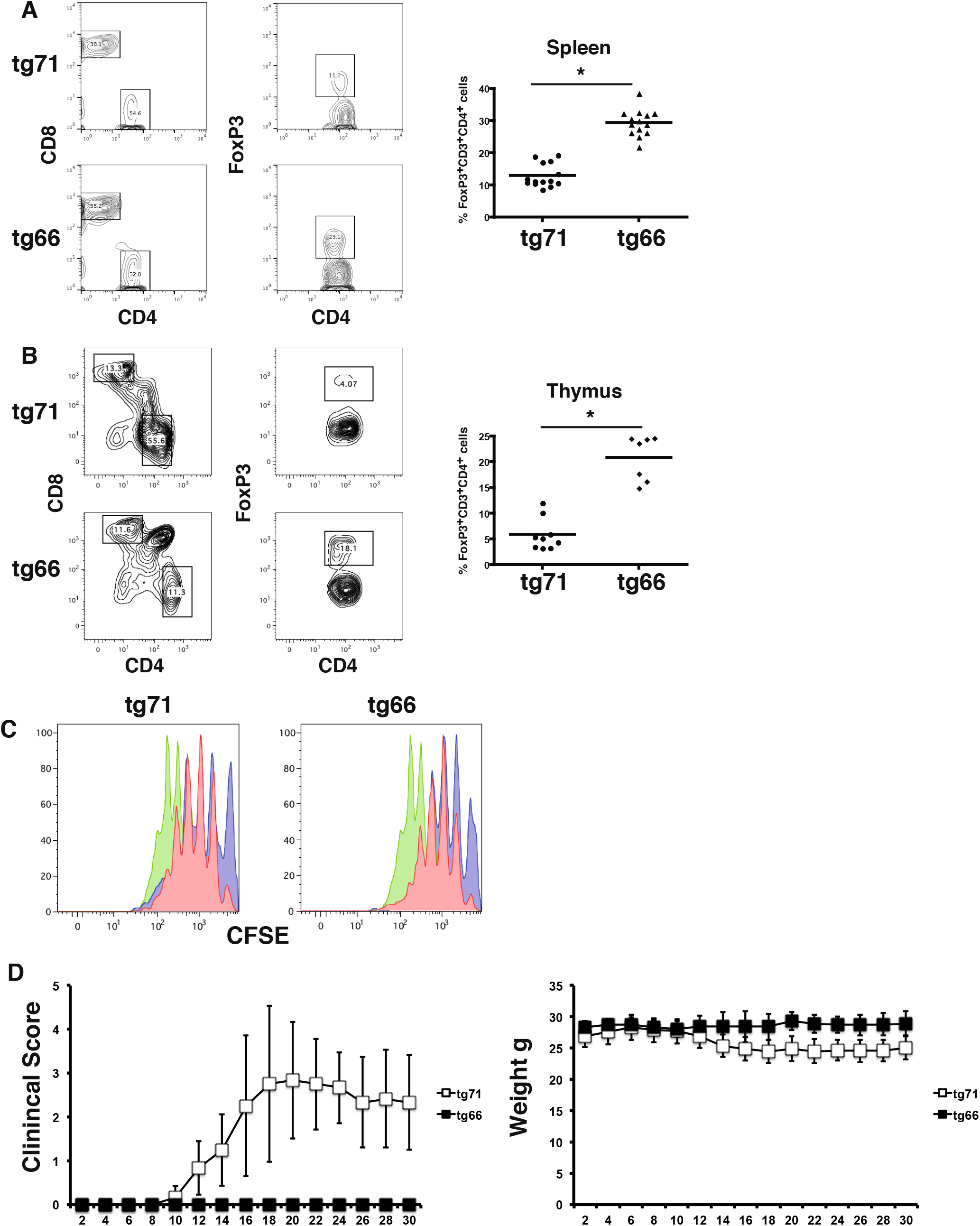
Increased frequency of Tregs in tg66 mice. (A) Frequency of Foxp3^+^ T cells in the spleens of tg66 mice (p<0.05 Mann-Whitney test). (B) Frequency of Foxp3^+^ T cells among CD3^+^CD4^+^ T cells in the thymus of tg66 mice. (C) T cell suppression assay demonstrating that Tregs from tg66 mice are equally capable of inhibiting stimulated CD8^+^ T cells as Tregs from tg71 mice. Green 0 Treg, Blue 2 Treg:1 CD8^+^ T cell, Red 1 Treg:2 CD8^+^ T cells. (D) Induction of EAE in tg66 mice is inhibited while tg71 mice develop disease and suffer weight loss.

Since Notch receptors have also been previously shown to be protective in autoimmune diseases such as EAE (27) and diabetes (28), we injected both tg66 and tg71 mice with MOG in order to induce EAE. Interestingly, while weight loss and progressive development of motor disability for EAE was observed in tg71 mice, little or no symptoms were evident in tg66 mice during the 30-day observation period (Fig. 3D).

In addition to the increased frequency of Tregs within the thymic CD4^+^ T cell population of tg66 mice, the phenotype of these Tregs found in the tg66 mice was also different to Tregs found in the tg71 mice. Thymic Tregs in tg66 mice had an increased frequency of IL-33R (ST2) expressing cells compared to Tregs from tg71 mice (Fig. 4A). Expression of ST2 on Tregs is associated with more suppressive functionality of Tregs, associated with high expression levels of IL-10 and TGFβ (29). KLRG1 has been identified as another marker for terminal differentiation in Tregs (30, 31). In addition, CD103 expression on Tregs has also been associated with increased suppression capacity by Tregs (32, 33). When the frequencies of CD103^+^KLRG1^−^, CD103^+^KLRG1^+^ and CD103^−^KLRG1^+^ subsets of Foxp3^+^ Tregs were examined in thymic Tregs, there were significant increases in all three subsets in the tg66 mice when compared to tg71 mice (Fig. 4B). Similarly, ICOS expression on Tregs has also been associated with suppressive activity of Treg by aiding IL10 production but has also been shown to play a role in Treg survival (34). Thymic Tregs from tg66 mice exhibited increased expression of ICOS on their surface when compared to their counterparts from the tg71 mice. Finally, the glucocorticoid-induced tumour necrosis factor receptor-related protein (GITR) expression on Treg has been associated as marker for activated Tregs (35) and GITR expression was significantly higher on Tregs from tg66 compared to tg71 thymic Tregs (Fig. 4C and D). Overall, these data demonstrate that the Tregs from tg66 mice have a phenotype of mature, highly activated and highly immunosuppressive Tregs.

**Figure 4.**
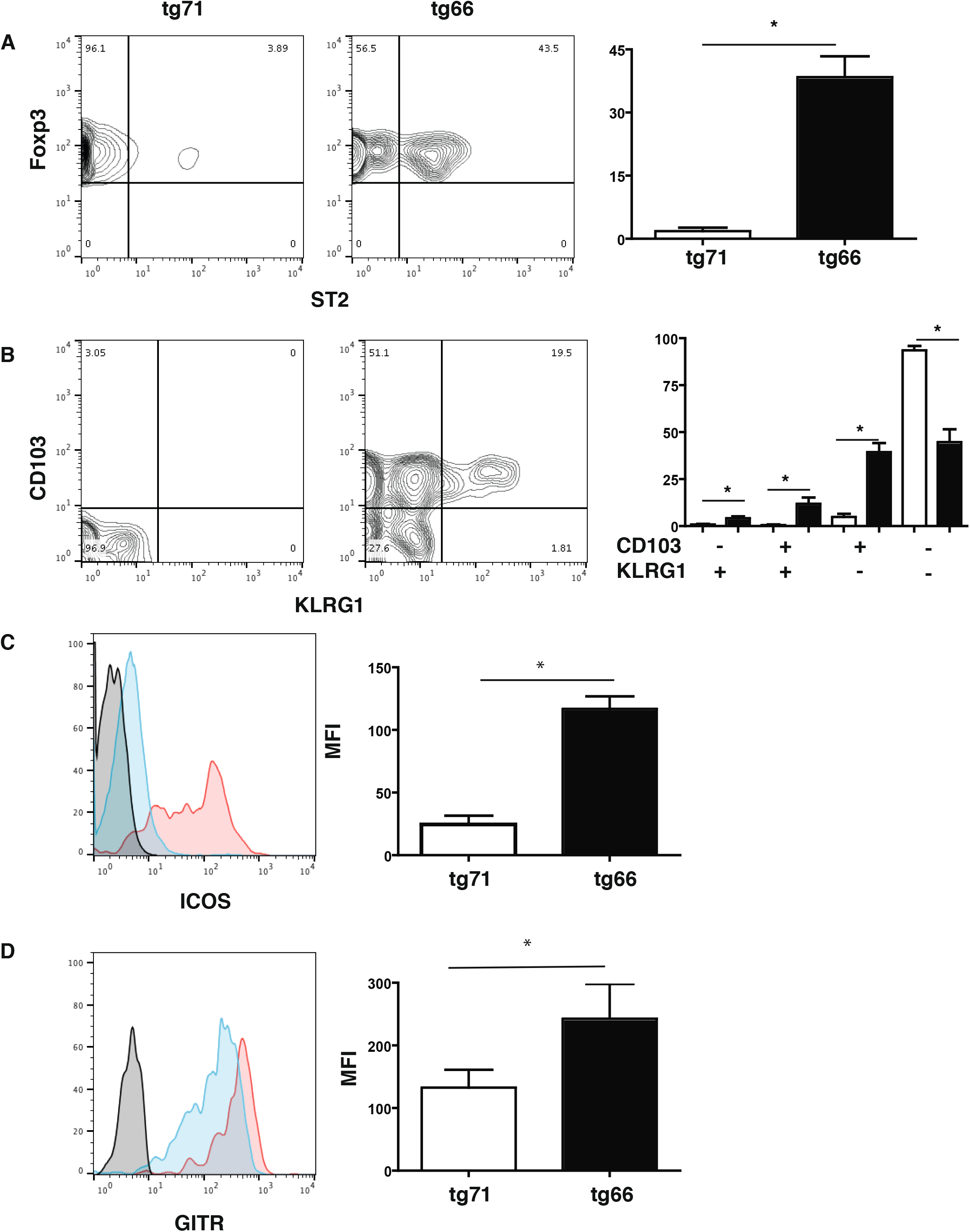
Phenotype of Tregs from tg66 mice. (A) Frequency of ST2 (IL-33R) on thymic Tregs from tg66 mice. (*p<0.05 Mann Whitney test 5-8 mice) (B) Frequency of CD103 and KLRG1 on thymic Tregs from tg66 and tg71 mice. Bar graph represents data from n=6-8 from 3 experiments. (*p<0.05 Mann Whitney test n=5-8 mice) (C) Expression of ICOS and (D) GITR on thymic Foxp3^+^ Tregs from tg66 (solid line) and tg71 mice (dashed line). (*p<0.05 Mann Whitney test n=5-8 mice)

### Tregs from tg66 mice expressed increased levels of granzyme B and C

Notch and Notch ligand signaling plays a role in granzyme and perforin induction (36, 37), which could induce cell-mediated killing of TECs. Examining the CD8 T cells from the thymus, we found that they expressed more EOMES, perforin and granzyme B than CD8 T cells from tg71 mice (Supplemental Figure 2). Expression of granzymes and perforin has also been reported in Tregs (38). Examining the T cells of the 14 day-old thymi, the Foxp3^+^CD4^+^ SP Tregs from tg66 mice had higher frequencies of granzyme B and perforin when compared to those from tg71mice (Fig. 5A and C). In addition, in tg66 mice nearly all Tregs expressed granzyme C, while in tg71 mice this proportion was only 25% of Tregs (Fig. 5B). Tg66 mice were crossed to *pfp*^−/−^ mice to determine whether thymic atrophy was associated with perforin. However, this was not the case as the T cells numbers were still reduced tg66 x *pfp*^−/−^ mice (Fig. 5D).

**Figure 5.**
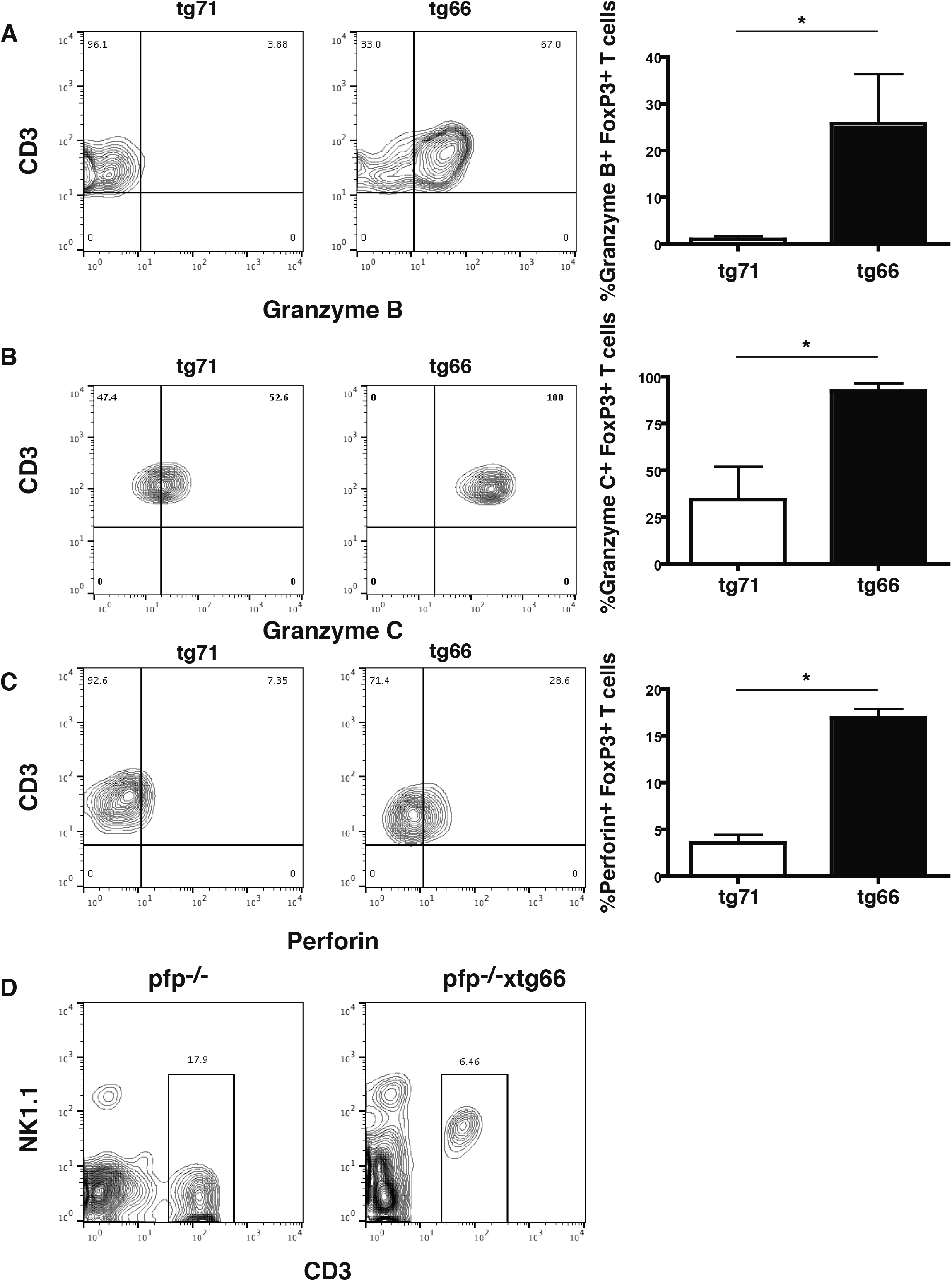
Increased expression of granzymes and perforin in thymic Tregs from tg66 mice. (A) Frequency of granzyme B-expressing thymic Tregs from tg71 and tg66 mice. (B) Frequency of granzyme C-expressing thymic Tregs from tg71 and tg66 mice. (C) Frequency of perforin-expressing thymic Tregs from tg71 and tg66 mice. *p<0.05 Mann Whitney test n=6-8. (D) Tg66 crossed with *pfp*^−/−^ mice still have reduced T cell output in the spleen: *pfp*^−/−^ (left) and tg66 x *pfp*^−/−^ (right) mouse.

### Tg66 bone marrow chimeras have reduced thymocyte numbers

To test if Jagged1 over-expression could affect T cell development in an already formed thymus, bone marrow from tg66 and tg71 mice were transplanted into cγRag^−/−^ mice that lack lymphocytes and so have reduced thymus size. While in the periphery, we found no differences in the frequency of T cells populations between mice receiving either bone marrow from tg66 or tg71 mice (Fig6 A and B), in mice receiving tg66 bone marrow the thymi were still relatively smaller, on average we only recovered around 6×10^5^ thymocytes from mice receiving tg66 bone marrow, compared to 7×10^6^ observed in chimeric mice receiving tg71 bone marrow (Fig. 6C). The frequency of thymic Tregs among the CD4^+^ T cells was still significantly increased in the thymi of the mice receiving tg66 bone marrow (Fig. 6D and E). However, when mice were treated with anti-Treg depleting or anti-Jagged1 blocking antibody there was 5-6-fold increase in the number of thymocytes (approximately 3×10^6^), which was related to a concomitant reduction in the number of Tregs (Fig. 6C and E). These data demonstrate that Jagged1 was driving Treg development in the thymus and that these Treg modulated thymocytes.

**Figure 6.**
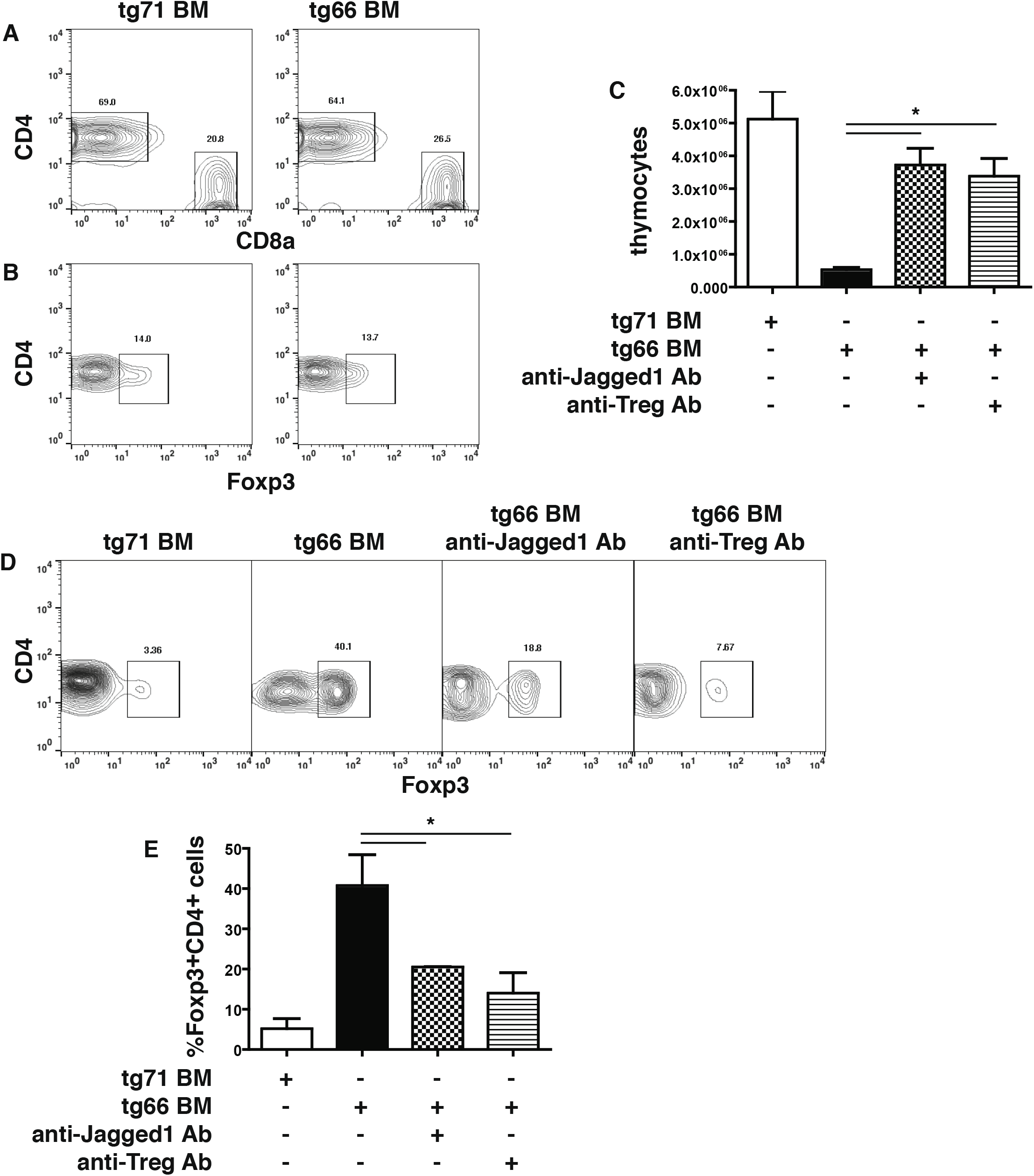
Reduced thymic size is related to increased frequency of Tregs in chimeric mice and can be reversed by blocking Jagged1. (A) Reduced thymic size in chimeric mice receiving tg66 BM. Thymic size increased upon blocking Jagged1 or depleting Tregs with anti-CD25 antibody (b and c) Frequency of thymic Tregs is increased in chimeric mice receiving tg66 bone marrow. The frequency of thymic Tregs is reduced by treating the mice with anti-Jagged1 or anti-Treg Ab.

## Discussion

In the tg66 mouse strain we determined that Jagged1 expression on T cells was increased tenfold and that this led to a complete loss of the thymus at day 28. Jagged1 expression on pre-T cells could, either through cis- or trans-interaction with Notch receptors on their surface, induce changes in the CD4/CD8 ratio, leading to the skewing of the tg66 T cells towards CD8^+^ T cells. Furthermore, among the reduced CD4^+^ T cells the frequency of Foxp3^+^ T cells was significantly increased in tg66 mice. These Tregs appeared to function normally *in vitro* since they could inhibit CD8^+^ T cell proliferation following CD3/CD28 activation. Furthermore, the Tregs in tg66 mice had a more distinct phenotype with increased expression levels of CD103, KLRG1 and ST2, when compared to tg71 mice. Finally using bone marrow chimera mice we found that these Tregs could inhibit the development of thymocytes but this was blocked by anti-Jagged1 antibodies.

The Notch system contains four receptors (Notch1-4) and two sets of ligands (Delta (1, 3 and 4) and Jagged (1 and 2). Both Delta and Jagged proteins activate Notch through proteolytic cleavage in the transmembrane domain of Notch by a γ-secretase (39) and γ-secretase inhibitors (GSI) potently prevent activation of Notch (39). Cleavage by γ-secretase releases the intracellular domain of Notch (NICD), which then translocates to the nucleus and binds to the DNA-binding protein RBPJ. The NICD and RBPJk then form a complex with Mastermind-like family (MAML) proteins and assemble a transcriptional activator complex. This trimolecular complex then recruits additional co-activators, including chromatin-modifying enzymes, to activate the transcription of Notch target genes. Notch can also signal through two non-canonical pathways, one through the mTOR/AKT pathway and the other through NFκB activation (40).

Notch expression on T cells plays a major role in the T cell differentiation process as well as in determination of the different T cell subsets (4, 26, 41). Lack of Notch1 prevents T cell development and instead leads to intrathymic development of B cells. Similar findings were also made in *RBJ*-deficient mice, since RBJ mediates signaling of all Notch receptors. The thymic stroma expresses all the ligands for Notch, suggesting that these signals are important for T cell development. Furthermore, Jagged1 is expressed on T cell subsets in the periphery (42). However, the mechanisms as to how Notch signaling induces different T cell subsets is still unclear, with different results obtained depending on the models used as well as differences from *in vivo* and *in vitro* observations (4).

Increased numbers of Tregs induced by Jagged1 in the thymus might lead to thymic aplasia since TGFβ affects thymic development (43). Several studies have associated Treg development with Notch signaling, noting that Notch signaling can enhance Treg differentiation and function *in vitro* (41). Notch1 and TGF-β signaling pathways cooperatively regulate Foxp3 expression and Treg maintenance both *in vitro* and *in vivo* (24). Thus, Jagged1 expression on T cells may drive Treg development, leading to increased intrathymic TGFβ that in turn could lead to thymic loss.

The breakdown of the thymic architecture and induction of cell death in the thymus of tg66 mice did not appear to be mediated by the perforin/granzyme pathway, even though granzyme B and perforin in Tregs have been previously reported to effectively maintain immune responses (38). However, granzymes themselves can also target other proteins that would be essential for maintaining normal thymic development (44). Granzyme B can enzymatically cleave extracellular matrix proteins such as fibronectin, vitronectin and laminin (45). The increased levels of granzyme B produced by Tregs could break down the ordered structure of the thymus, thus causing apoptosis in the thymic epithelium. In addition, granzyme B also can catalytically activate IL-1α (46) and IL-18 (47). This suggests that within the thymus there may exist a more pro-inflammatory environment that potentially drives cellular death and thymic involution. Furthermore, granzyme B can break down β-glycans, leading to the release of TGFβ (48), which as stated above can also lead to thymic involution. Finally, granzyme B also cleaves Notch receptors intracellularly thereby effectively activating them (49). Thus, even though Jagged1 may drive differentiation of CD8^+^ T cells and skewing towards Tregs in the CD4^+^ compartment, as well as a concomitant increase in granzyme B in these cells, there is also the possibility that granzyme B may feedback and affect Notch signaling. Although the frequency of T cells expressing granzyme C was high, the function of granzyme C is still unclear since it is not as efficient in inducing cell death as granzyme B (50). While granzyme B cleaves after aspartic acid, granzyme C cleaves after aromatic amino acids and can induce caspase-independent cell death (50). Release of granzymes independent of perforin may thus still induce cell death of TECs that leads to the observed thymic loss.

Thymic loss in tg66 mice therefore appears to potentially be a multifactorial process induced by Jagged1. Combinatorial effects of TGFβ and granzymes could affect the thymic epithelium by undermining its architecture. While there has not been any association between Tregs and other diseases characterized by thymic aplasia, such as DiGeorge’s syndrome, in Down Syndrome there is an increase in Treg frequency and reduced thymic size. Additionally, Jagged1 expression is increased in hematopoietic stem cells in Down Syndrome (51, 52). Interestingly, Jagged1 expression levels in hematopoietic stem cells from children with Down Syndrome were equivalent to those observed in elderly controls (52). It is interesting to speculate if increased expression of Jagged1 on hematopoietic stem cells in Down Syndrome and the elderly might lead to increased expression of Jagged1 on developing T cells in the thymus and thereby to thymic involution. Alagille syndrome is a heritable form of childhood chronic liver disease associated with a mutation in *JAG1*. No studies have defined any defects in thymic development in Alagille syndrome, however, although there does appear to be a defect in the development of T cell responses and in particular of Th1 immune responses (53, 54).

It is still unclear why the thymus disappears during normal development in all mammals and almost all vertebrates. However, there obviously seems to exist a yet undetermined selective pressure despite the negative effects on adaptive immunity. Besides energy conservations during puberty and reproductive age, prevention of either hematological cancers or autoimmune diseases in older organisms have been proposed as evolutionary pressures leading to thymic shrinkage. Our data does not rule out any of these hypotheses. However, the fact that the overexpression of Jagged1, together with the resulting increase of FoxP3^+^ Tregs were resistant to EAE induction, may indicate that protection of autoimmune diseases is at least a reasonable possibility.

Understanding how Jagged1 plays a role in regulating T cells has two clinical benefits. It does appear that Jagged1 might be an important molecule in human autoimmune diseases (55) and in tumor biology (56). Jagged1 did appear to induce granzymes and EOMES in CD8^+^ T cells, which might be utilized to generate cytotoxic T cells with greater killing abilities. Furthermore, Jagged1 expressed on tumors might convert intra-tumoral T cells to regulatory T cells. Therefore, understanding how these responses can be manipulated could ultimately help generate more effective anti-tumoral T cells.

## Abbreviations

B6: C57BL/6
DLL: Delta-like
EOMES: Eomesodermin
MOG: myelin oligodendrocyte
EAE: experimental autoimmune encephalomyelitis
Treg: regulatory T cell

## Acknowledgements

This work was funded by the Swedish Cancer Society, Swedish Research Council, the Karolinska Institute Foundations, and the Swedish Foundation for Strategic Research.

