## Supplemental Figures for "Jagged1 overexpression on T cells induces thymic regulatory T cells leading to thymic involution"

Running Title: Jagged1 and thymic involution

Joanna S. Kritikou[1,2], Irene Sánchez-Pascual[1], Juan Pedro Muñoz-Miranda[1,3], Neha Vashist[1,4], Arnika K. Wagner[2,5], Xing-Mei Zhang[5], Erika Assarsson[1], Margarita Dominguez-Villar [|6], Hideo Yagita[7], Francisco Garcia-Cozar[3], Peggy Riese[4], Robert A. Harris[5], Hans-Gustaf Ljunggren[1], Benedict J. Chambers[1].

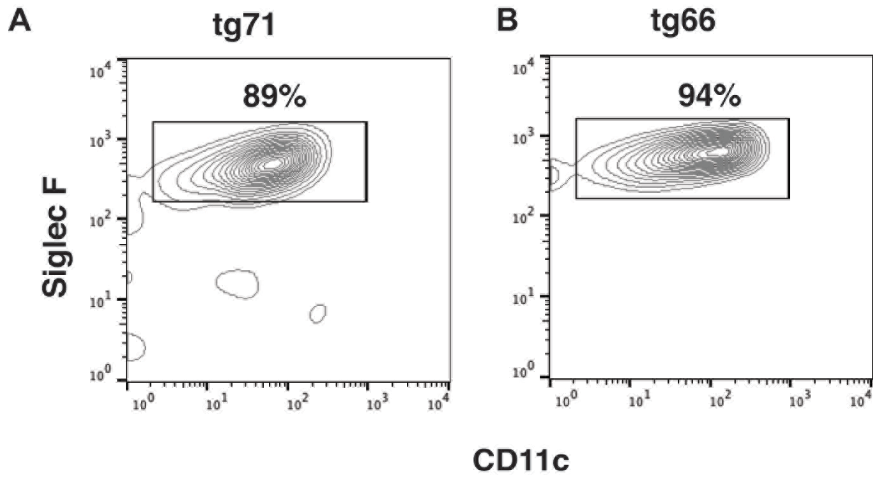


Supplemental Figure 1. Expression of Siglec F on granulocytes obtained from (A) tg71 and (B) Jagged1 overexpressing tg66 mice.

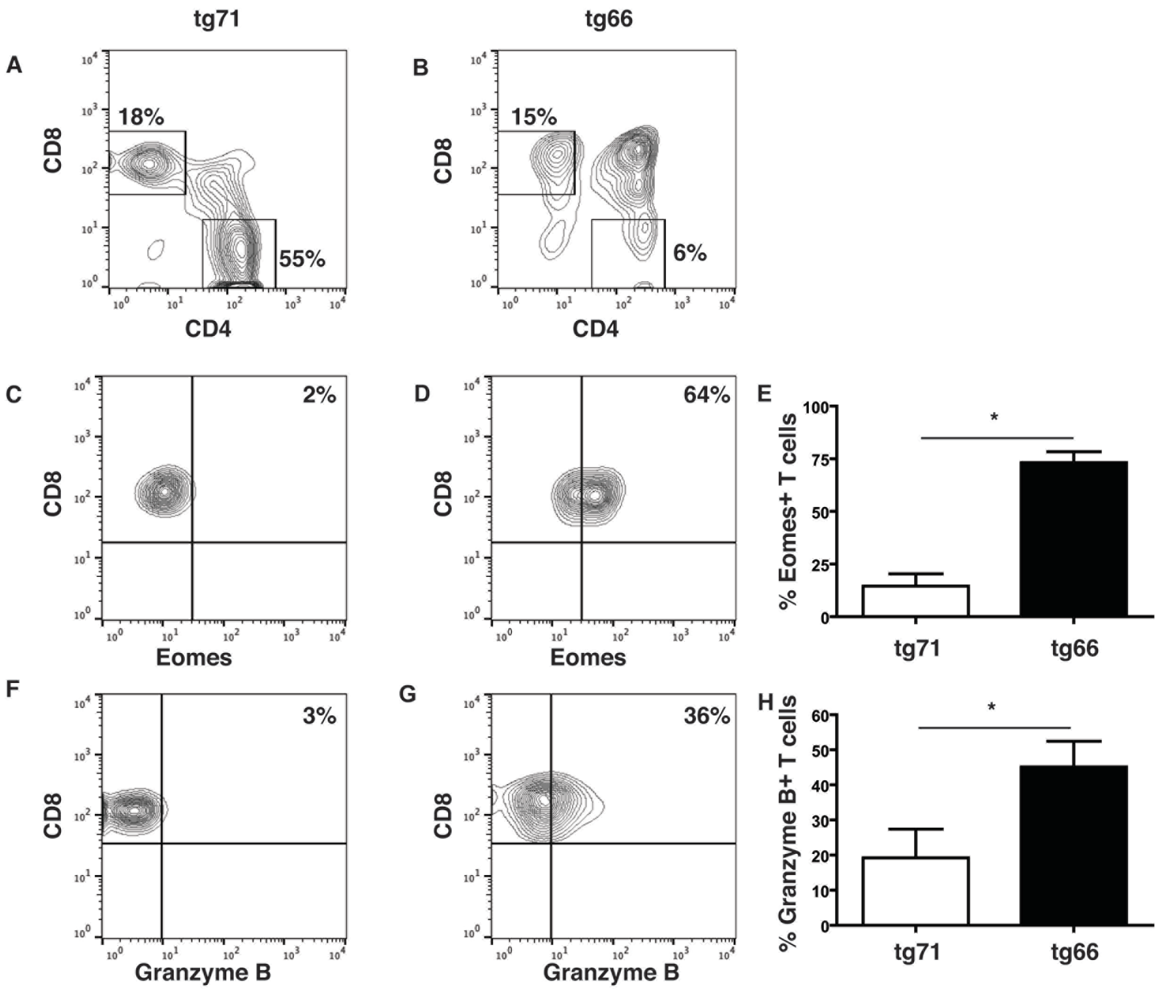


Supplemental Figure 2. CD8+ T cells from Jagged1 overexpressing tg66 mice express increased levels of EOMES and granzyme B. (A and B) By gating on CD3+ T cells in the thymus a decreased frequency of CD4 T cells is observed in tg66 mice. (C) Expression levels of the EOMES in tg71 and (D) tg66 mice. (e) Bar graph represents the percentage of EOMES-expressing T cells in tg71 and tg66 mice ($p<0.05$ Mann Whitney test, n=6-8). (F) Expression levels of granzyme B in tg71 and (G) tg66 mice. (H) Bar graph represents the percentage of granzyme B-expressing T cells in tg71 and tg66 mice ($p<0.05$ Mann Whitney test, n=6-8).
